# Within-group disagreements delay collective movement decisions in wild birds and primates

**DOI:** 10.64898/2026.11.19.767964

**Authors:** Damien R. Farine

## Abstract

One of the benefits of shared decision-making is that it can resolve conflicting preferences among group members. This allows the members of moving animal groups to maintain cohesion while deciding on where to go next. However, followers are faced with decisions that vary in complexity. At some times followers only have a single initiator to choose from, while at other times they may have to select among multiple simultaneous initiators each moving in different directions. One question is whether increased disagreement, in terms of the number and spread of directions among initiators, has an impact on the time it takes for followers to make their move. Here I use whole-group GPS tracking datasets from vulturine guineafowl (*Acryllium vulturinum*) and olive baboons (*Papio anubis*) to link the complexity of the decision arising from disagreement to the time it takes for followers to resolve a movement decision. Across both species, there is a significant impact of having more initiating directions, with every additional direction adding approximately 11 s (for 1 direction) to 99 s (for 9 directions) on decision time. This suggests that both suffer from the paradox of choice. In vulturine guineafowl, high directional conflict also added up to 162 s on decision times relative to low direction conflict, whereas baboons appeared unaffected by directional conflict. Overall, these results confirm that within-group disagreement among initiators can increase the complexity of decisions faced by followers, and that this ultimately increases the time needed to resolve movement decisions.

## INTRODUCTION

One of the challenges of living in a group with stable membership is that group members must reconcile differences in their preferences and decide about where to go (Byrne 2000) and what to do next (Kao *et al*. 2024). It is now well understood that most social animal species rely on shared decision-making (Strandburg-Peshkin *et al*. 2018). In shared decisions, any group member can contribute towards a possible outcome (e.g. by attempting to lead, or ‘initiating’ movement (King 2010; Liao *et al*. 2024)) and the group resolves the decision by selecting the outcome with the most support (typically following a ‘majority rule’ (Conradt & Roper 2005; Conradt & Roper 2007)). However, reaching a resolution takes time, which can be costly. For example, animals face time pressures to fit all their necessary activities in a day (Dunbar, Korstjens & Lehmann 2009; Gavrilets, Auerbach & van Vugt 2016), while slowing down or stopping reduces the energetic efficiency of movements (Klarevas-Irby, Nyaguthii & Farine 2025). In theory, the more conflict there is within a group, such as the more varied the directions of the initiators, the longer it should take to resolve the decision (Figure 1). However, the impact of within-group disagreements on the time taken to resolve collective decisions has not been widely explored.

**Figure 1.**
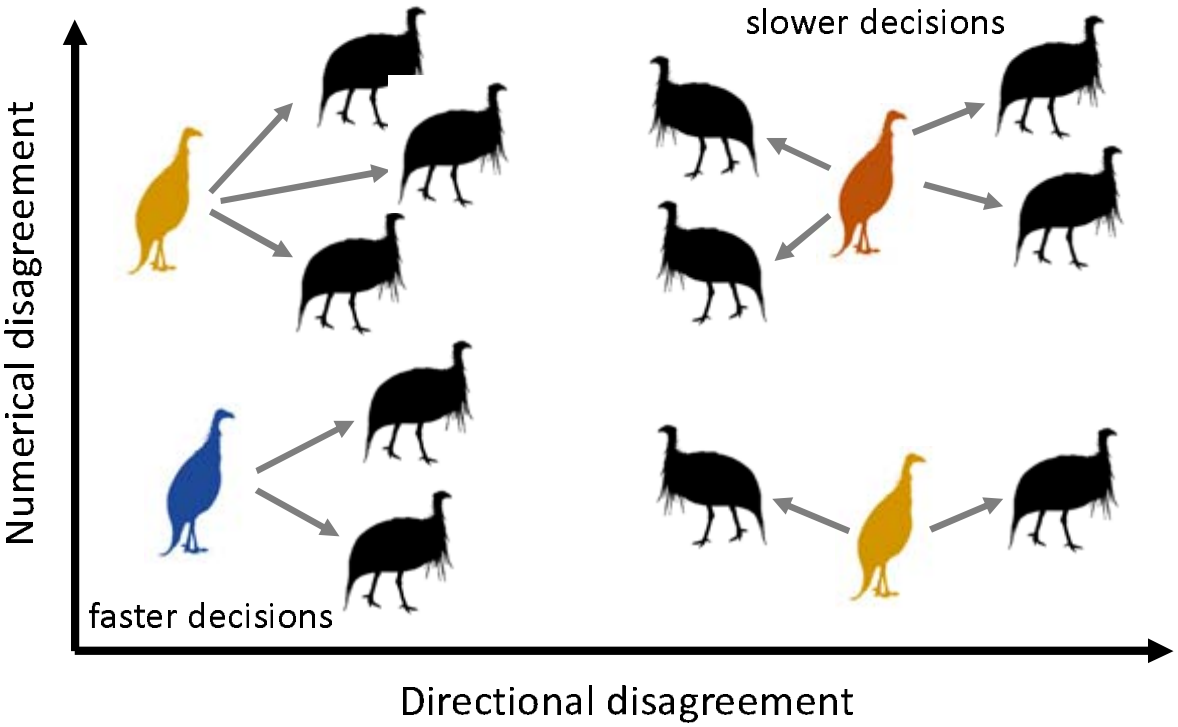
Greater disagreement within the group is predicted to increase the complexity of the decision, resulting in followers taking longer to decide which direction to take. Disagreement can be quantified along two axes. The directional disagreement, which is greater if the options are in more different directions, and numerical disagreement (i.e. the paradox of choice), which is greater when there are more options to choose from.

Having greater disagreement among group members should increase the complexity of reaching an outcome. In humans, people take longer to make choices when faced with more options (Hendrick, Mills & Kiesler 1968), which is known as the ‘paradox of choice’. For example, it can take longer to decide between products in a supermarket when there are more options to choose from. Similarly, if a single initiator in a moving animal group wants to go in a given direction, the decision can be easily (and rapidly) resolved, whereas reaching an outcome when several group members initiate in several and very different directions is more complex. To date, studies of collective decision-making have typically focused primarily on studying how groups resolve binary options (Chow, Rotstein & Garnier 2026). For example, theoretical studies have considered groups with two sets of preferences (Couzin *et al*. 2005), many lab studies have tracked groups using two-option y-mazes (Chow, Rotstein & Garnier 2026), and studies looking at collective decisions in wild animal groups have often simplified the analyses by focusing on cases with just two competing directional options (Strandburg-Peshkin *et al*. 2015; Papageorgiou, Nyaguthii & Farine 2024). Similarly, the difference in directions among initiators might also alter the time needed to reach consensus. For example, if two group members (or two clusters of group members) initiated in similar directions, then followers can simply select an average direction by moving in between the two directions (Couzin *et al*. 2005; Strandburg-Peshkin *et al*. 2015; Papageorgiou, Nyaguthii & Farine 2024). By contrast, if their initiations are 180 degrees apart, then followers must stop and must choose one direction versus the other (Couzin *et al*. 2005; Strandburg-Peshkin *et al*. 2015; Papageorgiou, Nyaguthii & Farine 2024).

Here I revisit data from two groups (three datasets) of wild vulturine guineafowl (*Acryllium vulturinum*) (Papageorgiou, Nyaguthii & Farine 2024; Brandl *et al*. 2025) and one group of wild olive baboons (*Papio Anubis*) (Strandburg-Peshkin *et al*. 2015) to test whether increased disagreement among group members results in delayed movements by followers. In all three groups, the majority of group members (~80%) were fitted with a GPS tag collecting data at 1 Hz (1 location per second). In both species, groups remain cohesive over the course of the day, meaning that all directional conflicts must be resolved before the group can move on. By analysing data on how group members moved relative to one another, the previous studies have found that group members resolve directional conflicts (when there were two clusters of concurrent initiators) using a majority rule. That is, they move in the direction of the largest cluster. However, these studies did not quantify the time taken for the follower to resolve conflicts and re-join the initiator(s), or how this is affected by the number of initiators and their directional differences. Here I explore all cases with one or more clusters of initiators (minimum two initiators), testing the prediction that situations with greater disagreement (more clusters and/or greater directional differences) are more complex to resolve, and thus take longer for followers to resolve.

## METHODS

### Data collection

I analysed published data from four whole-group GPS deployments fitted to vulturine guineafowl (three datasets) and olive baboons (one dataset). Data from both species were collected at the Mpala Research Centre in Laikipia District, Kenya outside of breeding seasons. The olive baboon dataset contained several independent juveniles whereas the vulturine guineafowl dataset was collected during a period where no juveniles were present. Details about the datasets can be found in Table 1.

**Table 1.** Summary of the leadership events data. Number of pulls is the number of cases where at least one initiator acted on a follower and where the follower responded to an initiator by following. Datasets 1 and 2 are from groups 1 and 2 in Papageorgiou *et al*. (2024) (Papageorgiou, Nyaguthii & Farine 2024). Dataset 3 is from Brandl *et al*. (2025) (Brandl *et al*. 2025). Dataset 4 is from Strandburg-Peshkin *et al*. (2015) (Strandburg-Peshkin *et al*. 2015). Note that group 1 in dataset 1 and dataset 3 are from the same group, collected 2 years apart.

| dataset | species | days of data | group size | # of pulls | 95% follow time range (s) | mean follows per id per day | mean daily follower time (mins) per id |
|---|---|---|---|---|---|---|---|
| 1 | Vulturine guineafowl | 83 | 26 | 56,174 | 23 – 876 | 38 | 146 |
| 2 | Vulturine guineafowl | 39 | 11 | 3,999 | 19 – 852 | 11 | 33 |
| 3 | Vulturine guineafowl | 50 | 20 | 27,609 | 30 – 788 | 42 | 159 |
| 4 | Olive baboon | 14 | 25 | 19,730 | 21 – 806 | 76 | 296 |

### Extracting decision-making events

The previous studies implemented the pull-anchor algorithm from Strandburg-Peshkin *et al*. (2015) to identify initiators and followers. This algorithm works by extracting sequences of minima to maxima to minima dyadic distances among group members, determining which of the two individuals contributed more to the increase in distance (labelled the initiator), and classifying the outcome based on who contributed to the subsequent decrease (pull if this was caused by the other individual moving, or anchor if this was caused by the original initiator). Specifically, initiation events were detected from periods where dyadic distance initially increased (from t1 to t2) and then decreased (from t2 to t3), with thresholds applied for minimum movement distances (a minimum change in distance of 3.5 m for vulturine guineafowl and 5 m for olive baboons). The initiator was defined as the individual contributing most to the increase in distance from t1 to t2. The outcome of the decision was then determined by the relative contributions from t2 to t3. If the initiator moved most, then the event was coded as an anchor (or failed initiation). If the other individual moved most, then the event was coded as a pull (successful initiation). In this paper, I only used data from successful initiations (pulls).

The pull-anchor algorithm operates at the dyadic level. To quantify events at the group level, the initiations were clustered in time using a chain rule, such that all events acting on the same follower that consecutively overlapped in time were considered as part of the same event. Within these events, the individual initiators were clustered into sets of co-initiators using a circular Gaussian mixture model. The number of clusters was recorded, as well as their directional disagreement. Disagreement was quantified using the circular variance, giving values between 0 (no disagreement, all initiators are in the same direction) and 1 (high disagreement, initiators are uniformly distributed across possible directions). Circular variance was calculated based on the directional vectors from the follower to each initiator, hence accounting for potential differences in the number of initiators across clusters.

Both Strandburg-Peshkin *et al*. (Strandburg-Peshkin *et al*. 2015) and Papageorgiou *et al*. (Papageorgiou, Nyaguthii & Farine 2024) restricted their analyses to cases where there were two clusters of initiators, as their focus was on which direction followers would select when deciding. Here I examine the impact of the number of clusters as well as the directional disagreement (higher disagreement is when initiators are more equally distributed across 360°) among clusters on the time taken for a follower to re-join the successful initiator (or cluster of initiators) in successful pulls. Because longer chains would naturally add both more clusters and a longer time, I recorded the follower time as starting when the maximum dyadic distance was reached with the successful initiator (or cluster) and finishing when the follower has rejoined the initiator or cluster of initiators (the subsequent minimum distance). This serves as a measure of follower decision-making time that is independent of the timing of the movements by unsuccessful initiators.

### Data analysis

I extracted the number of clusters and the directional disagreement for each event from each dataset. I then calculated the follower time, which starts when the follower first starts reducing the distance to the initiator and finishes when the follower has re-joined the initiator. I defined this as the decision-making time of the follower (noting that I included only successful pull events). I then built an identical generalised linear model (using the lmerTest (Kuznetsova, Brockhoff & Christensen 2017) library in R) for each dataset. This model included follower time as a response variable and number of clusters and disagreement as predictor variables. I log-transformed the follower time and square-root-transformed disagreement to improve model fit, but these transformations did not qualitatively impact the results. I also fitted follower ID as a random effect.

## RESULTS

Great disagreement among initiators leads to significantly longer decision-making time by followers (Figures 2-3; Table 2). In all three vulturine guineafowl dataset and in the baboon dataset, it took longer for followers to join initiators when there were more clusters of initiators. In vulturine guineafowl, having greater directional disagreement also increases decision-making time, such that followers took longer when clusters were more equally distributed around possible directions. Despite the difference in terms of the effect of directional agreement between guineafowl and baboons, the patterns are remarkably consistent across all four datasets. Specifically, they have similar ranges of decision-making times (Table 1, intercepts in Table 2) and similar effects of number of clusters (coefficients in Table 2). Further, estimates from the same social group (datasets 1 in Figure 2A and dataset 3 in Figure 2C) are very similar despite being collected 2 years apart.

**Table 2.** Summary of model results. Different models were fitted to each of the four datasets, allowing direct comparisons between them. Significance values are given by stars (*: <0.05, **: <0.01, ***: <0.001), unless specified. Note that the response variable is log transformed. Time delay estimates given in the abstract were calculated using an intercept of 4.8, a slope for number of clusters of 0.07 (with no directional disagreement), and a disagreement of 0.85 (for vulturine guineafowl only).

|  | Dataset 1 |  |  | Dataset 2 |  |  | Dataset 3 |  |  | Dataset 4 |  |  |
|---|---|---|---|---|---|---|---|---|---|---|---|---|
|  | β±SE | t | P | β±SE | t | P | β±SE | t | P | β±SE | t | P |
| Intercept | 4.76±0.01 | 364 | *** | 4.71±0.01 | 76.4 | *** | 5.0±0.00 | 204 | *** | 4.86±0.02 | 150 | *** |
| Disagreement | 0.76±0.02 | 33.5 | *** | 0.44±0.01 | 5.2 | *** | 1.3±0.05 | 19.3 | *** | -0.55±0.03 | -0.75 | n.s. |
| # of clusters | 0.06±0.00 | 31.1 | *** | 0.11±0.02 | 8.8 | *** | 0.06±0.00 | 24.6 | *** | 0.07±0.00 | 22.1 | *** |

**Figure 2.**
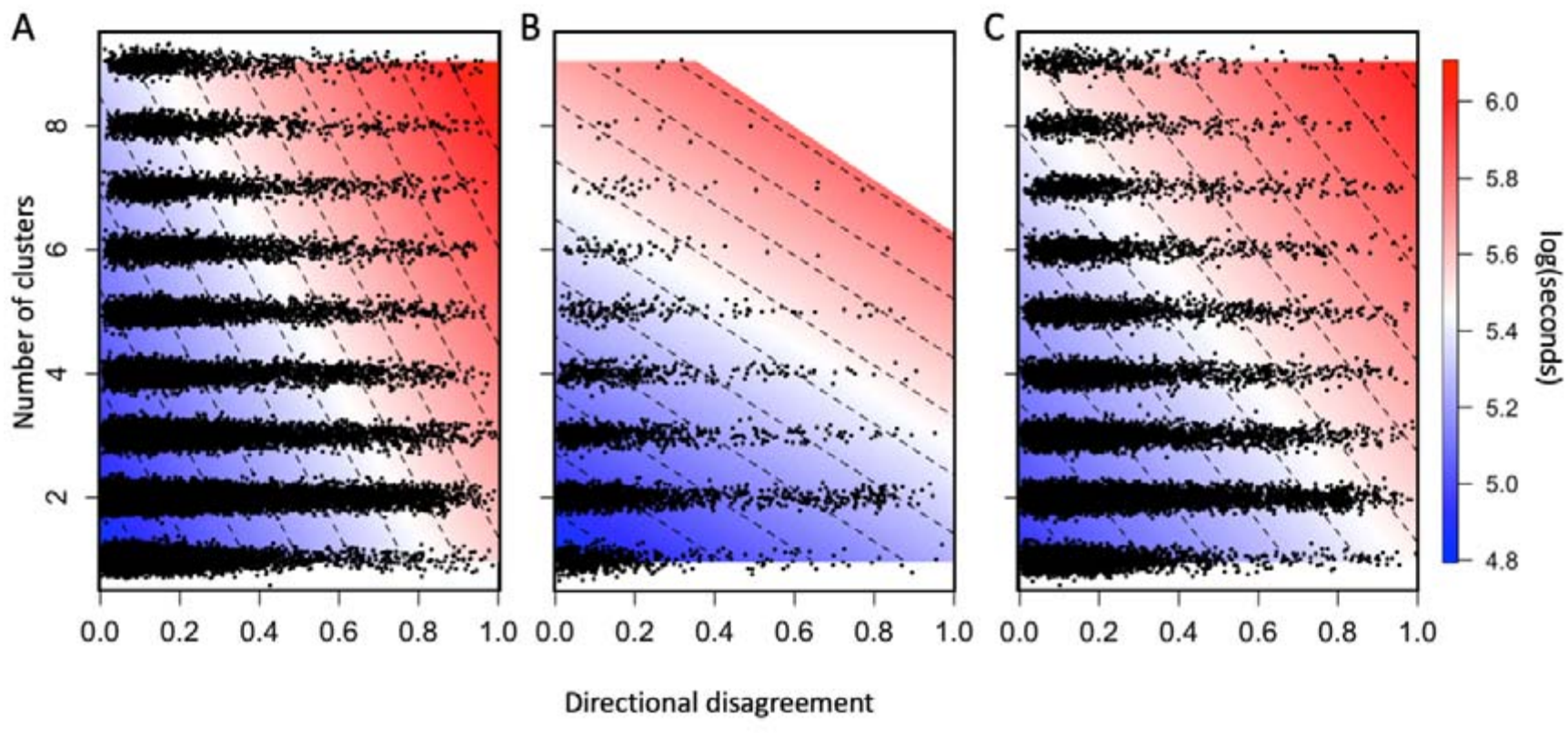
Increasing numerical and directional conflicts result in longer delays by followers in vulturine guineafowl. Each panel shows data from one dataset (A–C correspond to datasets 1–3 in Table 1, respectively). As both the directional disagreement and the number of distinct clusters of initiators increase, the time taken by followers (the natural log of time in seconds, shown in the background colour) increases. Low disagreement is when initiators are all moving in the same direction, whereas high disagreement represents cases where initiators are equally spread across all possible directions (e.g. 180° in cases with two intiators). Colours and dashed lines show the model predictions, while points show the spread of the raw data across the two axes of disagreement (i.e. showing the predictor variables only). Model results are given in Table 2.

**Figure 3.**
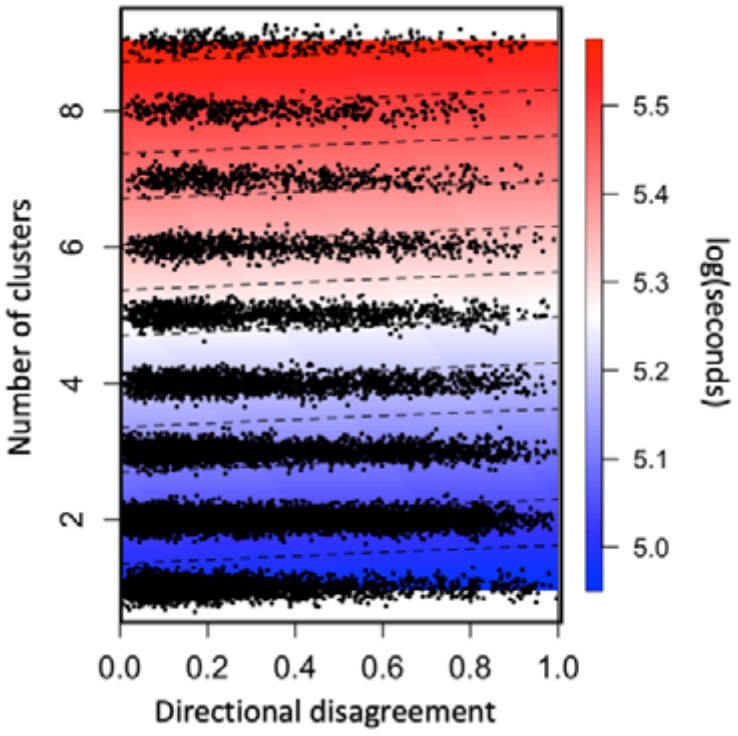
Increasing numerical conflicts result in longer delays by followers in olive baboons. Figure components are as per Figure 2 (but for dataset 4 in Table 1). Model results are given in Table 2.

## DISCUSSION

Collective decision-making takes time (Gavrilets, Auerbach & van Vugt 2016), and more complex decisions are hypothesised to require longer to resolve (Chow, Rotstein & Garnier 2026). Here I showed that increased disagreement among initiating baboons and vulturine guineafowl had longer delays by followers. Specifically, followers took longer to re-join initiators when there were more simultaneous clusters of initiators and (in guineafowl) when the directional disagreement among simultaneous initiators higher. These results support the predictions from theory (Chow, Rotstein & Garnier 2026). They also suggest that disagreement among initiators can be a cost of living in groups with stable membership by causing delays to movement decisions. These delays can subsequently reduce movement efficiency (Klarevas-Irby, Nyaguthii & Farine 2025) and reduce time available for other activities for both followers (who are deciding) and initiators (who are waiting).

Surprisingly few studies have investigated the time associated with reaching consensus in collective decision-making. Among these, two studies from Trinidian guppies (*Poecilia reticulata*) highlight both the challenge associated with reaching a decision and the drive to do so efficiently. The first study showed that the challenge of reaching consensus is consistent across group sizes. Specifically, while larger groups were faster to reach the fork (decision-point) of a y-maze, groups of all sizes took a consistent amount of time to decide between which fork to take (Wade, Ramnarine & Ioannou 2020). Simulations also suggest that group size should have little effect on the speed of decision-making beyond very small groups (Ward *et al*. 2008). By contrast, a study conducted in giraffes (*Giraffa sp*.) found that larger groups take longer to resolve decisions, and ultimately were more likely to split (Kasozi *et al*. 2026). In the present study, I found consistent effects across all group sizes (see parameter estimates in Table 2), but that individuals in the larger group of vulturine guineafowl and in olive baboons spent also took substantially more time to resolve conflicts among initiators. The second study showed that guppies became faster at making decisions over time when repeatedly faced with the same y-maze (Hansen *et al*. 2021). This highlights a general increase in efficiency of decision-making with greater familiarity of the environment. However, this increase can only reach a certain point, and I expect that both the baboon and guineafowl groups will have reached this point long before the studies took place.

One question is what drives directional disagreement among group members. Previous modelling work (Conradt & Roper 2003; Davis, Crofoot & Farine 2022; Farine, Davis & Crofoot 2026) suggests that differences in time budgets can introduce conflicts in terms of the timing of decisions (i.e. when to move), and it is well established how animals can use quorum rules to overcome these costs (Conradt & Roper 2005; Sumpter & Pratt 2009; Farine 2022; Kao *et al*. 2024). However, what has received less attention is the potential role of environmental complexity. Environments or contexts with more options, such as more foraging areas, could increase disagreement among group members. While quantifying the targets of collective movements in free-ranging animals is challenging, there is some evidence for the effect of environmental complexity on decision-making in movements by herds of Przewalski horses (*Equus ferus przewalskii*). A previous study found that herds take longer to make collective movements when these movements spanned multiple habitat types (Bourjade *et al*. 2009). Hence, greater disagreement—and hence decision-making complexity—could be linked to situations when the environment naturally raises more options. This highlights a likely strong feedback between spatial features of the environment and social processes within the group (Cantor *et al*. 2021; Webber *et al*. 2023).

While the effect of disagreement among group members is relatively consistent among vulturine guineafowl and olive baboons, there may be some species-specific differences. In both species, followers took longer to decide when there were more options, demonstrating that individuals across these species suffer from the paradox of choice. However, olive baboon followers did not appear to be affected by the amount of directional disagreement. This could be because they are faced with such conditions much more often. Despite both living in the same location, and thus moving through the same habitat, 13.5% of decisions by the baboon group had a disagreement of greater than 0.5, whereas fewer than 5% of vulturine guineafowl decisions had such high levels of disagreement. This difference likely arises due to the costs of disagreements, with vulturine guineafowl being much more susceptible to predation and thus maintaining very high levels of social cohesion (Betti *et al*. 2026). Comparisons of leadership data also revealed that initiating vulturine guineafowl were much more likely to be successful than initiating olive baboons (Papageorgiou, Nyaguthii & Farine 2024), even though both species initiators were less likely to be followed (i.e. events were more likely to result in an anchor) when initiators had greater directional disagreement (Strandburg-Peshkin *et al*. 2015; Papageorgiou, Nyaguthii & Farine 2024). The lack of effect of disagreement on the following time by baboons could thus reflect some adaptations to resolving such conflicts, either via their cognitive ability or other mechanisms that are yet to be discovered.

While the current study addresses one key gap—namely the effect of disagreement on the time it takes for followers to follow, and thus groups to reach consensus when making movement decisions—there remains several important questions. One of these is whether followers and groups have strategies to reduce the complexity associated with high disagreements. In social insects, colonies can efficiently select among options (e.g. new nest sites) by evaluating options concurrently (Franks *et al*. 2003). However, this option is not as clearly available for organisms on the move. Two key mechanisms could help simplify decision-making in moving animal groups. One is quorum decision-making, and specifically a sub-majority rule, whereby the group moves when some threshold of individuals have made a decision. This could be the case in vulturine guineafowl, which will make collective departures once 13 individuals have initiated movement (Papageorgiou & Farine 2020). A second is to turn the decision into multiple smaller decisions. Laboratory studies suggest that animals can reduce complexity by breaking the problem up into a series of binary movement decisions (Sridhar *et al*. 2021). Hence, there remains open questions about how moving animal groups overcome the costs of collective decision-making.

A further challenge faced by animals making collective movements is that the geometry of groups (and thus of the decision) does not remain fixed. Every movement (e.g. one individual following one initiator) changes the relative balance of options and directions, both for itself and for any of its potential followers. This could both increase complexity (if followers constantly have to ‘start afresh’) or decrease it (if sets of followers cumulatively reduce the number of options simultaneously by reinforcing a subset of these). More research is needed on the spatiotemporal dynamics of groups making collective movement decisions, most notably on how the geometry of the decision-making landscape changes over time.

Increasing disagreement among group members slows down follower behaviour. This insight aligns with concepts of increasing complexity of options (more options) resulting in slower collective decision-making. Such an outcome is likely to also have important ecological consequences for followers (who may take longer to make decisions), for initiators (who may have to wait for followers), and for the group as a whole. For example, speed affects travel time between patches, which has strong impacts on the dynamics of leadership and the amount of conflict in terms of preferences among group members (Farine, Davis & Crofoot 2026). Furthermore, future work should investigate whether individuals maintain other activities during these conflicts (e.g. continuing foraging), which can be achieved by tracking activity using accelerometers (Christensen *et al*. 2025). This will be important to determine if collective decisions indeed induce missed time for other activities and if the delaying decisions induces additional opportunity costs (alongside losses in movement efficiency). Either way, these results add to a growing literature revealing that while collective behaviours can bring benefits to animal groups, they also introduce novel challenges and costs.

## ACKNOWLEDGEMENTS & FUNDING

This research was funded by the European Research Council (ERC) under the European Union’s Horizon 2020 research and innovation program (grant agreement number 850859).

## REFERENCES

Betti, L., Musciotto, F., Papageorgiou, D., Battiston, F. & Farine, D.R. (2026) Beyond the dyad: higher-order structure within cohesive animal groups can reveal novel patterns of sub-group dynamics. Animal Behaviour, in press.

Bourjade, M., Thierry, B., Maumy, M. & Petit, O. (2009) Decision-Making in Przewalski Horses (Equus ferus przewalskii) is Driven by the Ecological Contexts of Collective Movements. Ethology, 115, 321–330.

Brandl, H.B., Klarevas-Irby, J.A., Zuñiga, D., Wheat, C.H., Christensen, C., Omengo, F., Nzomo, C., Cherono, W., Nyaguthii, B. & Farine, D.R. (2025) The physiological cost of leadership in collective movements. Current Biology, 35, 4003–4010.

Byrne, R.W. (2000) How monkeys find their way. Leadership, coordination and cognitive maps of African baboons. On the move: how and why animals travel in groups (eds S. Boinski & P.A. Garber). University of Chicago Press, Chicago, IL.

Cantor, M., Maldonado-Chaparro, A.A., Beck, K.B., Brandl, H.B., Carter, G.G., He, P., Hillemann, F., Klarevas-Irby, J.A., Ogino, M., Papageorgiou, D., Prox, L. & Farine, D.R. (2021) The importance of individual-to-society feedbacks in animal ecology and evolution. J Anim Ecol, 90, 27–44.

Chow, P.C.K., Rotstein, H.G. & Garnier, S. (2026) How do Cockroach Groups Integrate Multiple Attributes in a Best-of-N Task? bioRxiv, 10.64898/62026.64807.64824.740418.

Christensen, C., Ferreira, A.C., Cherono, W., Maximiadi, M., Nyaguthii, B., Ogino, M., Herrara, D.C. & Farine, D.R. (2025) Moving towards more holistic machine learning-based approaches for classification problems in animal studies. Animal Behaviour, 230, 123386.

Conradt, L. & Roper, T.J. (2003) Group decision-making in animals. Nature, 421, 155–158.

Conradt, L. & Roper, T.J. (2005) Consensus decision making in animals. Trends in Ecology & Evolution, 20, 449–456.

Conradt, L. & Roper, T.J. (2007) Democracy in animals: the evolution of shared group decisions. Proceedings of the Royal Society B-Biological Sciences, 274, 2317–2326.

Couzin, I.D., Krause, J., Franks, N.R. & Levin, S.A. (2005) Effective leadership and decision-making in animal groups on the move. Nature, 433, 513–516.

Davis, G.H., Crofoot, M.C. & Farine, D.R. (2022) Using optimal foraging theory to infer how groups make collective decisions. Trends in Ecology & Evolution, 37, 942–952.

Dunbar, R.I.M., Korstjens, A.H. & Lehmann, J. (2009) Time as an ecological constraint. Biological Reviews, 84, 413–429.

Farine, D.R. (2022) Collective action in birds. Current Biology, 32, R1140–R1144.

Farine, D.R., Davis, G.H. & Crofoot, M.C. (2026) Optimal foraging can drive emergent initiator-follower dynamics in social groups. Oikos, 2026.

Franks, N.R., Mallon, E.B., Bray, H.E., Hamilton, M.J. & Mischler, T.C. (2003) Strategies for choosing between alternatives with different attributes: exemplified by house-hunting ants. Animal Behaviour, 65, 215–223.

Gavrilets, S., Auerbach, J. & van Vugt, M. (2016) Convergence to consensus in heterogeneous groups and the emergence of informal leadership. Scientific Reports, 6.

Hansen, M.J., Burns, A.L., Monk, C.T., Schutz, C., Lizier, J.T., Ramnarine, I., Ward, A.J.W. & Krause, J. (2021) The effect of predation risk on group behaviour and information flow during repeated collective decisions. Animal Behaviour, 173, 215–239.

Hendrick, C., Mills, J. & Kiesler, C.A. (1968) Decision time as a function of the number and complexity of equally attractive alternatives. Journal of Personality and Social Psychology, 8, 313–318.

Kao, A.B., Banerjee, S.C., Francisco, F.A. & Berdahl, A.M. (2024) Timing decisions as the next frontier for collective intelligence. Trends in Ecology & Evolution, 39, 904–912.

Kasozi, H., Mutebi, M., Muge, J., Kakuru, A., Strandburg-Peshkin, A. & Briefer, E.F. (2026) The socioecological correlates of collective movements in giraffes. Behavioural Ecology, 37, arag098.

King, A.J. (2010) Follow me! I’m a leader if you do; I’m a failed initiator if you don’t? Behavioural Processes, 84, 671–674.

Klarevas-Irby, J.A., Nyaguthii, B. & Farine, D.R. (2025) Moving as a group imposes constraints on the energetic efficiency of movement. Proceedings of the Royal Society B-Biological Sciences, 292.

Kuznetsova, A., Brockhoff, P.B. & Christensen, R.H.B. (2017) lmerTest Package: Tests in Linear Mixed Effects Models. Journal of Statistical Software, 82, 1–26.

Liao, C.C., Magrath, R.D., Manser, M.B. & Farine, D.R. (2024) The relative contribution of acoustic signals versus movement cues in group coordination and collective decision-making. Philosophical Transactions of the Royal Society B-Biological Sciences, 379.

Papageorgiou, D. & Farine, D.R. (2020) Shared decision-making allows subordinates to lead when dominants monopolize resources. Science Advances, 6.

Papageorgiou, D., Nyaguthii, B. & Farine, D.R. (2024) Compromise or choose: shared movement decisions in wild vulturine guineafowl. Communications Biology, 7.

Sridhar, V.H., Li, L., Gorbonos, D., Nagy, M., Schell, B.R., Sorochkin, T., Gov, N.S. & Couzin, I.D. (2021) The geometry of decision-making in individuals and collectives. Proceedings of the National Academy of Sciences of the United States of America, 118.

Strandburg-Peshkin, A., Farine, D.R., Couzin, I.D. & Crofoot, M.C. (2015) Shared decision-making drives collective movement in wild baboons. Science, 348, 1358–1361.

Strandburg-Peshkin, A., Papageorgiou, D., Crofoot, M.C. & Farine, D.R. (2018) Inferring influence and leadership in moving animal groups. Philosophical Transactions of the Royal Society B-Biological Sciences, 373.

Sumpter, D.J.T. & Pratt, S.C. (2009) Quorum responses and consensus decision making. Philosophical Transactions of the Royal Society B-Biological Sciences, 364, 743–753.

Wade, A.S.I., Ramnarine, I.W. & Ioannou, C.C. (2020) The effect of group size on the speed of decision making depends on compromise and predation risk across populations in the guppy. Behaviour, 157, 1173–1192.

Ward, A.J.W., Sumpter, D.J.T., Couzin, L.D., Hart, P.J.B. & Krause, J. (2008) Quorum decision-making facilitates information transfer in fish shoals. Proceedings of the National Academy of Sciences of the United States of America, 105, 6948–6953.

Webber, Q.M.R., Albery, G.F., Farine, D.R., Pinter-Wollman, N., Sharma, N., Spiegel, O., Vander Wal, E. & Manlove, K. (2023) Behavioural ecology at the spatial-social interface. Biol Rev Camb Philos Soc, 98, 868–886.

